# *Zfp423*, a Joubert syndrome gene, is a domain-specific regulator of cell cycle progression, DNA damage response and Purkinje cell development in the cerebellar primordium

**DOI:** 10.1101/139691

**Authors:** Filippo Casonil, Laura Crocil, Camilla Bosonel, Roberta D’Ambrosio, Aurora Badaloni, Davide Gaudesi, Valeria Barilil, Justyna R. Sarna, Lino Tessarollo, Ottavio Cremona, Richard Hawkes, Søren Warming, G. Giacomo Consalez

## Abstract

Neurogenesis is a tightly regulated process whose success depends on the ability to balance the expansion/maintenance of an undifferentiated neural progenitor pool with the precisely timed birth of sequential generations of neurons. The *Zfp423* gene encodes a 30-Zn-finger transcription factor (TF) that acts as a scaffold in the assembly of complex transcriptional and cellular machineries regulating neural development. While null mutants for *Zfp423* feature a severe cerebellar hypoplasia, the underlying mechanism is only partially characterized. Mutations of the human ortholog *ZNF423* have been identified in patients carrying cerebellar vermis hypoplasia (CVH) or Joubert Syndrome (JS), associated with other signs of classical ciliopathy outside the central nervous system (CNS). ZNF423 also plays a role in the DNA damage response (DDR). To further characterize the role of ZFP423 in cerebellar neurogenesis, with a focus on Purkinje cells (PC) development, we analyzed two previously undescribed mutant mouse lines carrying allelic in-frame deletions of the corresponding gene, selectively affecting two functionally characterized protein-protein interaction domains, affecting zinc (Zn) fingers 9-20 or 28-30. Some phenotypic defects are allele specific: *Zfp423^Δ9-20/Δ9-20^* mutants exhibit a depletion of the OLIG2+ PC progenitor pool in the cerebellar ventricular zone (VZ). In these mutants, M-phase progenitors display changes in spindle orientation indicative of a precocious switch from symmetric to asymmetric cell division. Conversely, the *Zfp423^Δ28-30/Δ28-30^* primordium displays a sharp decrease in the expression of PC differentiation markers, including CORL2, despite an abundance of cycling PC progenitors. Moreover, and importantly, in both mutants VZ progenitor cell cycle progression is remarkably affected, and factors involved in the DDR are substantially upregulated in the VZ and in postmitotic precursors alike. Our in vivo evidence sheds light on the domain-specific roles played by ZFP423 in different aspects of PC progenitor development, and at the same time supports the emerging notion that an impaired DNA damage response may be a key factor in the pathogenesis of JS and other ciliopathies.

## INTRODUCTION

Within the cerebellar primordium, two neurogenetic matrices are discernible: the ventricular zone (VZ) and the anterior rhombic lip (RL) (reviewed in Leto et al., 2015). Starting at E10.5–E11.5 (Goldowitz and Hamre, 1998), the cerebellar VZ, positive for PTF1A (Hoshino et al., 2005), neurogenin 1 and 2 (Salsano et al., 2007; Zordan et al., 2008) and ASCL1 (Grimaldi et al., 2009), and adhesion molecules NEPH3 and E-cadherin (Mizuhara et al., 2009) gives rise in overlapping waves to all γ-aminobutyric acid (GABA)-positive neurons, including PCs (Florio et al., 2012; Kim et al., 2011; Lundell et al., 2009) and GABAergic interneurons, which are born sequentially from a common PAX2+ progenitor pool (Leto et al., 2009; Leto et al., 2006; Maricich and Herrup, 1999; Weisheit et al., 2006).

Defects affecting the VZ or RL translate into congenital malformations of the cerebellum. Human congenital ataxias feature motor disability, muscular hypotonia and incoordination. Malformations involve the cerebellar vermis, and are sometimes accompanied by other neural and extraneural abnormalities. Joubert syndrome (JS, MIM21330) is a debilitating multisystem ciliopathy featuring a distinctive malformation of the brainstem and cerebellar peduncles called the molar tooth sign for its appearance in axial MRI sections. It is characterized by congenital malformation of the brainstem and agenesis or hypoplasia of the cerebellar vermis (CVH), leading to breathing dysregulation, nystagmus, hypotonia, ataxia, and a delay in achieving motor milestones, and possibly affecting cognitive function. JS is usually inherited as an autosomal recessive trait and is usually caused by mutations in genes encoding proteins that localize either in the primary cell cilium or centrosome / basal body / transition zone (reviewed in Romani et al., 2013). These proteins control ciliary structure / stability, and signal transduction (reviewed in Goetz and Anderson, 2010; Han and Alvarez-Buylla, 2010).

*Zfp423* encodes a 30-Zn-finger nuclear protein that works both as a scaffold and as a transcription factor. ZFP423 (or its human ortholog ZNF423) interacts with multiple regulatory molecules, including EBF transcription factors (Tsai and Reed, 1997; Tsai and Reed, 1998), involved in cerebellar development (Croci et al., 2011; Croci et al., 2006) and cortical patterning (Chung et al., 2008; Chung et al., 2009). ZFP423 also acts a coactivator in BMP (Hata et al., 2000), Notch (Masserdotti et al., 2010) and retinoic acid (Huang et al., 2009) signaling. *Zfp423*-deficient mice display a variety of developmental defects, including a completely penetrant cerebellar vermis hypoplasia (CVH) (Alcaraz et al., 2006; Cheng et al., 2007; Warming et al., 2006). Malformations of cerebellar hemispheres are dependent on modifier genes and stochastic factors (Alcaraz et al., 2011). Importantly, patients carrying *ZNF423* gene mutations / deletions have been diagnosed with JS, CVH, NPHP and other signs of ciliopathy (Chaki et al., 2012). While ZFP423 has been convincingly implicated in the cilium-mediated response to sonic hedgehog (SHH) during cerebellar granule cell (GC) proliferation (Hong and Hamilton, 2016), our observations clearly point to an additional key role for this protein in PC development long before the onset of GC clonal expansion. Incidentally, GC clonal expansion relies on SHH released by PCs starting around birth (Dahmane and Ruiz-i-Altaba, 1999; Wallace, 1999; Wechsler-Reya and Scott, 1999), so that the final number of GCs is heavily influenced by the total number of postmitotic PCs.

Importantly, ZFP423 / ZNF423 also interacts with Poly ADP-ribose polymerase 1 (PARP1) (Ku et al., 2006) and centrosomal protein 290 (CEP290) (Chaki et al., 2012). PARP1 is a double stranded (ds) DNA-damage sensor that recruits MRE11 and ataxia-telangectasia mutated (ATM) to sites of DNA damage, indirectly linking ZNF423 to the ATM pathway of DNA damage signaling. CEP290 is a centrosomal protein mutated in JS and NPHP, whose loss causes enhanced DNA damage signaling, DNA breaks, replication stress, and supernumerary centrioles (Slaats et al., 2015). More broadly, recent evidence also supports a role for ciliopathy genes in the DDR. Both CEP290 and NEK8 mutations lead to an accumulation of DNA damage due to disturbed replication forks (Choi et al., 2013; Slaats et al., 2015). Furthermore, increased DNA damage signaling has been detected in CEP164-, ZNF423-, and SDCCAG8-associated nephronophthisis (Airik et al., 2014; Chaki et al., 2012). Our findings indicate that the two allelic deletions have divergent effects on PC progenitor pool maintenance and PC progenitor differentiation. Conversely, both *Zfp423* deletions analyzed here cause a delay in PC progenitor cell cycle progression and a significant increase in the number of mitotic and postmitotic precursors featuring signs of DNA damage.

## MATERIALS AND METHODS

### Animal Care

All experiments described in this paper were performed in agreement with the stipulations of the San Raffaele Scientific Institute Animal Care and Use Committee (I.A.C.U.C.) and the Guide to the Care and Use of Experimental Animals from the Canadian Council for Animal Care.

### Mouse genetics

The *Zfp423* Δ9-20 and Δ28-30 mouse lines were generated by gene targeting. The targeting constructs were engineered using BAC recombineering in *E. coli*; the mutations were introduced into the *Zfp423* gene by homologous recombination in 129S1/SVImJ cells. The resulting mutants were then backcrossed repeatedly with C57BL/6N mice to generate congenic strains. All experiments were carried out on homozygous mutant animals starting from backcross generation N8, expected to carry 99,6% of the C57BL/6N genetic background. All studies were conducted in homozygous mutant embryos, using coisogenic wild type control littermates. Genotyping was done by PCR as described (Corradi et al., 2003) using allele-specific primers. Primer sequences are listed under Supplemental Materials and Methods.

### Dot Blot

P0 wild type and mutant cerebella were lysed in RIPA buffer containing protease inhibitors, and protein concentration was determined by the BCA assay (Pierce). Protein analysis was done using a dot-blot apparatus as previously described (Casoni et al., 2005). Briefly, polyvinylidene difluoride membrane (Millipore Corp.) was soaked in methanol for a few seconds and then in ultrapure water for 2 min and conditioned in 20 mM Tris-HCl, pH 7.4, prior to assembly in the apparatus (Bio-Dot Apparatus, Biorad). Deposition of each sample on the membrane was achieved by vacuum filtration. The membrane was probed with the rabbit polyclonal anti-Zfp423 antibody (1:1000, Santa Cruz). As a loading control, we used a mouse monoclonal anti-αTubulin antibody (1:3000, Sigma).

### Tissue Preparation

Pregnant dams were anesthetized with Avertin (Sigma) prior to cervical dislocation. Embryos were fixed 6-8 hours by immersion with 4% PFA in 1X PBS, cryoprotected overnight in 30% sucrose in 1X PBS, embedded in OCT (Bioptica), and stored at −80°C. For postnatal brains, mice were anesthetized with Avertin (Sigma), transcardially perfused with 0.9% NaCl, followed by 4% PFA in 1X PBS, and prepared for freezing as above. The brains and embryos were sectioned sagittally or frontally on a cryotome.

### EdU administration and labeling

Pregnant dams were injected intraperitoneally with 5-Ethynyl-2'- deoxyuridine (EdU, 50 μg/g body weight). Embryos were collected and treated as described above. EdU incorporation was revealed using the Click-iT^®^ EdU Cell Proliferation Assay kit (Life Technologies), according to manufacturer’s instructions.

### Immunofluorescence

Sections were washed in 1X PBS, blocked and permeabilized in 10% serum, 0.3% Triton X-100 in 1X PBS, and incubated with primary antibodies overnight at 4°C, rinsed, then incubated with secondary antibodies at room temperature for 2 hrs (1:1000 Alexa Fluor™, Molecular Probes®). Sections were counterstained with 4',6-diamidino-2-phenylindole (DAPI, 1:5000, Sigma) and mounted with fluorescent mounting medium (Dako). Peroxidase immunohistochemistry was performed as described in Sarna et al., 2006. Antibodies are listed under Supplemental Materials and Methods.

### In situ hybridization

Digoxigenin-labeled riboprobes were transcribed from plasmids containing *Lhx5* cDNAs. In situ hybridization was performed as described (Croci et al., 2011).

### Cell pair assay

E11.5 cerebellar primordia were dissected in HBSS, collected by centrifugation and dissociated by pipetting in culture medium containing DMEM (Sigma), Glutamax (Life Technologies), penicillin/streptomycin (Euroclone), 1mM sodium pyruvate (Life Technologies), 1mM N-acetyl-L-cysteine (Sigma), B27 and N2 (Life Technologies), and 10ng/ml FGF2 (Peprotech). Single cells were plated onto 24-well plates coated with poly-D-lysine (10μg/ml), at a density of 30000 cells/well. The cultures were maintained in a humidified incubator at 37°C with constant 5% CO_2_ supply. About 24 hrs later, the cultures were fixed and immunostained with anti-βIII-Tubulin (1:500, Covance) and anti-Sox2 (1:250, Abcam) Abs, and counterstained with DAPI (Sigma). Approximately 50 images per well were acquired using the GE Healthcare IN Cell Analyzer 1000 and analyzed with Adobe Photoshop.

### RT-qPCR

Total RNA was extracted with RNeasy MiniKit (Qiagen), according to manufacturer’s instructions. 1–1.5 μg of total RNA was retrotranscribed using first strand cDNA MMLV- Retrotranscriptase (Invitrogen) and random primers. Each cDNA was diluted 1:10, and 3 μl were used for each RT-qPCR reaction. mRNA quantitation was performed with SYBR Green I Master Mix (Roche) on a LightCycler480 instrument (Roche). Each gene was analyzed in triplicate, and each experiment was repeated at least three times. Data analysis was performed with the 2-ΔΔC(T) method. All RNA levels were normalized based upon *β-actin, H3f3a and Gapdh* transcript levels. For primer sequences see Supplementary Materials and Methods.

### Primer sequences

#### Genotyping primers

*Δ9-20*-wtF: 5’: GGCTTCCATGAGCAGTGCT-3’
*Δ9-20*-wtR: 5’: 3CTCCTGCAGGCTGTTGATGT-3’
*Δ9-20*-mutR: 5’: 3CGTCCACGTCATCCTCACT-3’
*Δ28-30*-wtF: 5’: 3TGAGAGAGGACACCTACTCT-3’
*Δ28-30*-wtR: 5’: GCAGGGAGCAAACTGTCTCTT-3’;
*Δ28-30*-mutR: 5’: GGTGTGACCTTTGTGCGAGA-3’

#### RTqPCR primers

*Zfp423* F: 5’: CTTCTCGCTGGCCTGGGATT-3’
*Zfp423* R: 5’: GGTCTGCCAGAGACTCGAAGT-3’
*β-actin* F: 5’: CTGTCGAGTCGCGTCCACC-3’
*β-actin* R: 5’: TCGTCATCCATGGCGAACTG-3’
*H3f3a* F: 5’: GGTGAAGAAACCTCATCGTTACAGGCCTGGTAC-3’
*H3f3a* R: 5’: CTGCAAAGCACCAATAGCTGCACTCTGGAAGC-3’
*Gapdh* F: 5’: TGAAGCAGGCATCTGAGGG-3’
*Gapdh* R: 5’: CGAAGGTGGAAGAGTGGGAG-3’

### Antibodies

Mouse monoclonal antibodies: Pancadherin (1:300, Sigma); βIII-Tubulin (1:500, Covance); Zebrin II (Brochu et al., 1990), used in spent hybridoma supernatant (1:5000); Phospho-histone H3 (1:1000, Abcam); Olig2 (1:200, Millipore). Rabbit antibodies included: Calbindin (1:1000, Swant); ZFP423/OAZ XL (1:1000, Santa Cruz Biotechnology); FoxP2 (1:1000, Abcam); Phospho-histone H3 (1:800, monoclonal, Millipore); Phospholipase Cß4 (PLCβ4) (1:1000, kind gift Dr. M. Watanabe, Hokkaido University, Japan); Sox2 (Abcam, 1:250); Olig2 (1:500, Millipore); Corl2 (1:2000, Aviva System Biology); γ-H2AX (1:200, Cell Signaling); 53BP1 (1:1000, Bethyl Lab). Goat antibody: LaminB (1:100, Santa Cruz). Signal from anti-OAZ antibodies was amplified using the Tyramide Signal Amplification Kit (Perkin Elmer), according to manufacturer’s instructions.

### Microscopy and image processing

Optical microscopy was carried out using a Leica SP8 microscope or a Zeiss AxioImager M2m microscope. Digital images were processed with Adobe Photoshop to adjust contrast and brightness according to Journal’s recommendations. Quantitative and morphometric evaluations were performed using the ImageJ software (NIH). In Figure 6C,D, signal was quantitated as the ratio of the area occupied by γH2AX to that occupied by DAPI (C), or EdU^3h^ (D). Quantitation was achieved with ImageJ using the selection tool to delimit areas of interest.

### Statistical analysis

Statistical analysis was performed in Graphpad Prism and was conducted as follows: a) the two-tailed unequal variance t test (Welch t-test) was employed to compare sets of quantitative values (wt vs. each mutant), b) categorical values were analyzed by means of the two-tailed Fisher’s exact test, and plotted as proportions, percentages or actual numbers, c) one-way ANOVA was calculated through the Kruskal-Wallis test followed by Dunn’s post hoc test for the comparison of areas occupied by different signals.

## RESULTS

### The ZFP423 protein is expressed throughout the VZ, peaking in M-phase ventricular zone progenitors

The distribution of the *Zfp423* transcript in the embryonic CNS has been described (Alcaraz et al., 2006; Cheng and Reed, 2007; Masserdotti et al., 2010; Warming et al., 2006). Here, we report the tissue and subcellular distribution of the corresponding protein in the early cerebellar primordium. On embryonic day 11.5 (E11.5), ZFP423 is abundantly expressed throughout the cerebellar primordium, especially flanking the midline. The protein is found in the rhombic lip (rl), in the roof plate (rp) (arrowheads in **A**), and in the prospective sub-arachnoid space (sa) (**Fig. 1A**). Both the rp and the sa space harbor signaling cells important for cerebellar development (inter alia, see Aldinger et al., 2009; Chizhikov et al., 2006). In the cerebellar VZ, which contains PC precursors, progenitor cell bodies known as radial glia (Anthony et al., 2004) are arranged into a pseudostratified epithelium and oscillate back and forth from the basal membrane to the apical margin (Florio et al., 2012), which contains an adherens junction belt (Kosodo et al., 2004). This process has been dubbed interkinetic nuclear migration (reviewed in Gotz and Huttner, 2005). The location of cell bodies correlates with specific phases of the cell cycle. At the apicalmost end of their range, bordering the ventricular lumen, progenitors enter M-phase and divide. In the cerebellar VZ, ZFP423 is expressed at its highest levels in M-phase precursors decorated by phosphorylated histone H3 (PHH3) (arrows in **Fig. 1B**). In the E12.5 VZ, robust ZFP423 expression levels are maintained in M-phase progenitors, and slightly lower levels are found in interphase nuclei scattered throughout the thickness of the VZ (arrows in **Fig. 1C**, magnification in **C’**). A subset of ZFP423 PC progenitors also expresses the marker OLIG2, mostly associated with G0/G1 PC progenitors (Ju et al., 2016). Likewise, the domain containing differentiating PC precursors positive for CORL2 colocalizes partially with the ZFP423+ postmitotic domain in the cerebellar primordium (**Supplemental Fig. 1**). These findings indicate that ZFP423 is expressed in PC progenitors.

**Figure 1.**
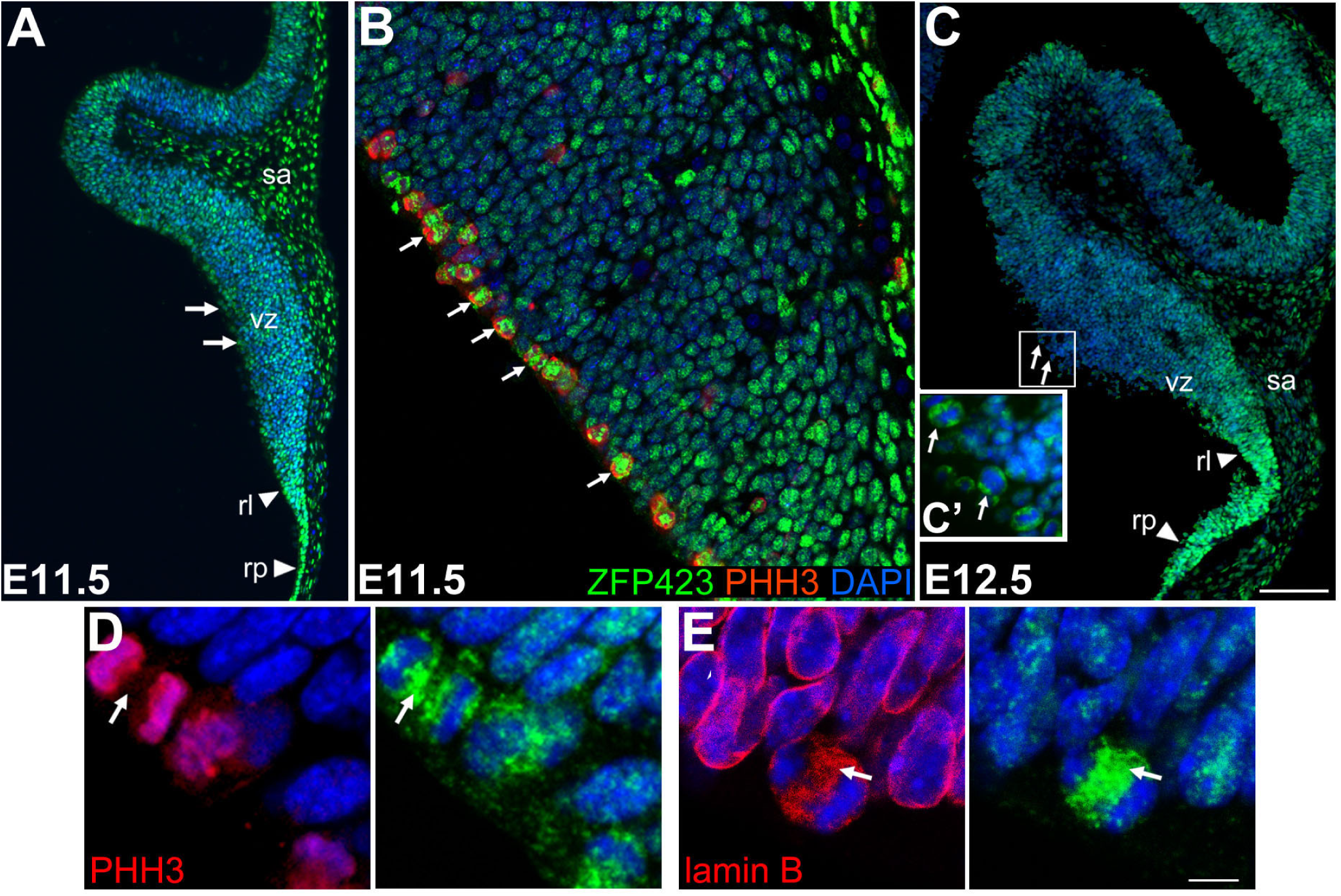
ZFP423 is expressed in both germinal layers of the cerebellar primordium. Immunofluorescence staining of cerebellar sagittal sections at E11.5 (**A, B, D, E**) and E12.5 (**C**). **A-C:** the ZFP423 protein is expressed at high levels in the cerebellar primordium, including the ventricular zone (arrows), the germinal layer which produces all GABAergic neurons and PCs in particular. **B:** ZFP423 is strongly expressed in radial glia progenitors positive for the M-phase marker PHH3 (arrows in **B**). **D, E:** whereas in interphase cells ZFP423 is mostly detected in the nucleus, in dividing PHH3+ progenitors the protein is mostly separate from chromatin (arrow in **D**). In some presumptive anaphase cells, ZFP423 colocalizes with the nuclear lamina protein lamin B (arrow in **E**). RL, rhombic lip; RP, roof plate; SA, subarachnoid space; VZ, ventricular zone. Size bar: 100μM (**A, C**); 50μM (**B**); 5μM (**D, E**).

At the subcellular level, most of the ZFP423 signal is uncoupled from PHH3+ chromatin (**Fig. 1D**) and overlaps partially with the nuclear intermediate filament lamin B (**Fig. 1E**). However, we have no conclusive *in vivo* evidence of ZFP423 colocalization with the centrosomal marker γ-tubulin (not shown, see Chaki et al., 2012).

### Generation and analysis of two allelic mutants harboring specific ZFP423 domain deletions

To dissect the domain-specific roles of ZFP423 in the events that lead up to PC neurogenesis, we produced two lines of mice by BAC recombineering in *E. coli* and gene targeting in ES cells. Each line carries a distinct targeted deletion of *Zfp423* (sketched in **Fig. 2A**). The first deletion, dubbed Δ9-20, spans nt 1258 through 2517 of the ORF (420 aa), encoding ZFs 9-20. The corresponding protein domain mediates a molecular interaction with the BMP-responsive element (BRE) and SMAD1-4 (Hata et al., 2000), and with PARP1 (Chaki et al., 2012; Ku et al., 2003). Moreover, this domain is required for the functional interaction with the Notch intracellular domain (Masserdotti et al., 2010). The second deletion, called Δ28-30, removes nt 3861-4108 of the ORF and causes a frameshift mutation that produces a stop codon 158 bp into exon 8. The resulting C-terminally truncated protein lacks ZFs 28-30 and 91 C-terminal aa. This domain has been implicated in the interaction with the Collier/Olf-1/EBF (COE) family of TFs (Tsai and Reed, 1997). Heterozygous mutants of both genotypes are viable and fertile, and display no overt phenotypic abnormality, suggesting that neither mutation has any relevant toxic gain of function or dominant negative effect (not shown). Conversely, homozygous mutants, on a C57BL/6N background, feature a highly significant increase in lethality compared to wt littermates by postnatal day 7 (χ^2^ test, *p*<0.0001). Survivors are profoundly ataxic (not shown). The analysis of young adult cerebella from outbred [(C57BL/6Nx129S1/SVImJ)F2] mutants (**Fig. 2B))** revealed that homozygotes for both mutations exhibit severe cerebellar hypoplasia, with a markedly reduced vermis. The Δ28-30 mutant features a pronounced posterior vermis deletion that is not observed in the Δ9-20. The corresponding mutant genes are upregulated with respect to the wt (**Fig. 2C)**). Both deleted proteins (sketched in **Supplemental Fig. 2**) retain the DNA-binding domain spanning ZFs 2-8 (Tsai and Reed, 1998) and are stable, as shown by protein dot blots of embryonic cerebellar lysates (**Fig. 2D)**). Deletion products retain a nuclear localization in E12.5 cerebellar primordia (**Fig. 2E)**). The results of these immunostainings were reproduced in cells transfected with fluorescently tagged constructs, ruling out non-specific antibody binding (not shown).

**Figure 2.**
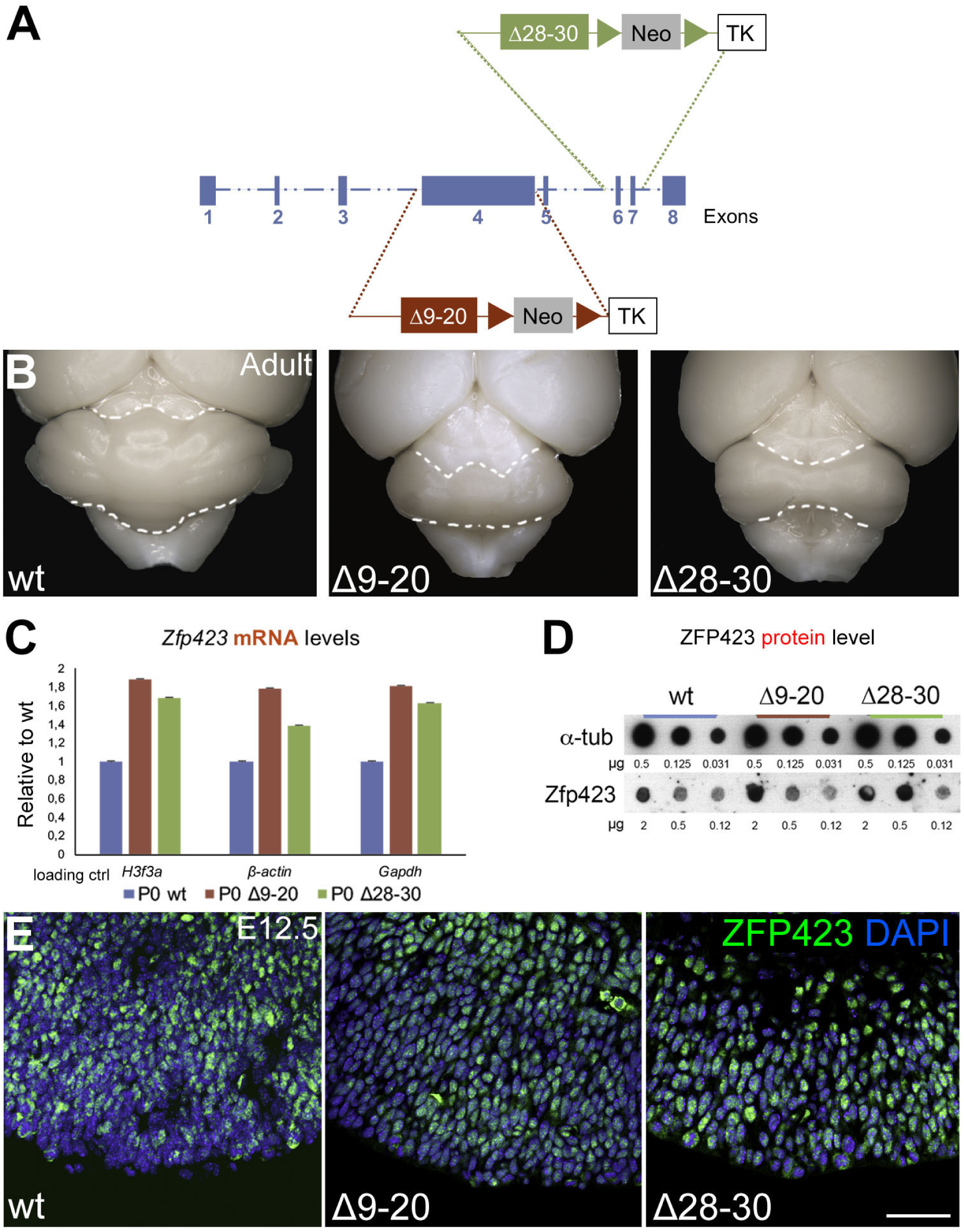
Both *Zfp423* deletion mutants produce a stable, predominantly nuclear protein and exhibit a severe cerebellar hypoplasia. **A:** Schematic representation of the constructs produced to generate, by homologous recombination, the Δ9-20 and Δ28-30 allelic mutants. **B**: dorsal view of cerebellar malformations in adult mutant mice, compared to the control. In homozygosity, both mutants display a decreased cerebellar size and a severe vermis hypoplasia, with a more pronounced posterior deletion in the Δ28-30. **C, D:** at P0, both deleted RNAs (RTqPCR in **C**) and proteins (dot blot in **D**) are expressed at robust levels. **E**: ZFP423 immunostaining of E12.5 sagittal cerebellar sections reveals that, *in vivo*, both deleted proteins are expressed and localize predominantly in the cell nucleus. Size bar: 25 μM.

### The number of PCs is decreased in both allelic *Zfp423* mutants

*Zfp423* null mutant embryos showed reduced BrdU incorporation in the cerebellar VZ at E14.5 (Alcaraz et al., 2006); however at that stage PC progenitors no longer proliferate. We conducted PC counts after the end of PC progenitor proliferation (E18.5, P17). These indicated a substantial drop in the number of PCs in both mutants. In particular, at P17, the perimeter of the PC layer in both mutants ranges between 27% and 60% of normal, depending on distance from the midline, with the sharpest decrease in the cerebellar vermis (**Fig. 3A)**), whereas the numerical density of calbindin+ cell bodies is normal in the mutant PC layer. These results demonstrate that *Zfp423* mutation causes a severe drop in PC number. In keeping with this evidence, the total PC number, as measured by counting immature FOXP2+ PCs in the fetal cerebellar cortex (E18.5), is sharply reduced in both allelic mutant cerebella (**Fig. 3C)**). Interestingly, the medial population depleted in the Δ9-20 cerebellum (**arrows in C**) corresponds to late-born PCs marked by birthdating with BrdU at 12.5 (**sketched in 3B**, Doorn and Hawkes, unpublished). Based on this evidence, we asked whether the PC loss scored in our allelic mutants affects both zebrinII + and zebrinII- PC subtypes or is selective for a specific subpopulation. In the adult cerebellum, PCs are arranged in two (mostly) mutually exclusive parasagittal patterns – one characterized by zebrin II/aldolase C expression (Brochu et al., 1990), the other by the expression of PLCβ4 (Sarna et al., 2006). These subsets correspond to early-born and late-born PC populations, respectively (Croci et al., 2006; Hashimoto and Mikoshiba, 2003; Larouche and Hawkes, 2006). Our results (**Fig. 3D)**) show that the Δ28-30 cerebellum, albeit smaller, contains a balanced complement of both PC types, while the Δ9-20 shows a selective loss of late-born (PLCβ4+) progenitors. Taken together, our findings suggest that both *Zfp423* mutants feature a severe loss of PCs, and that in the Δ9-20 cerebellar primordium PC progenitors may leave the cell cycle and differentiate prematurely, depleting the progenitor pool, in contrast to Δ28-30, where early- and late-born PCs are present in approximately normal proportions.

**Figure 3.**
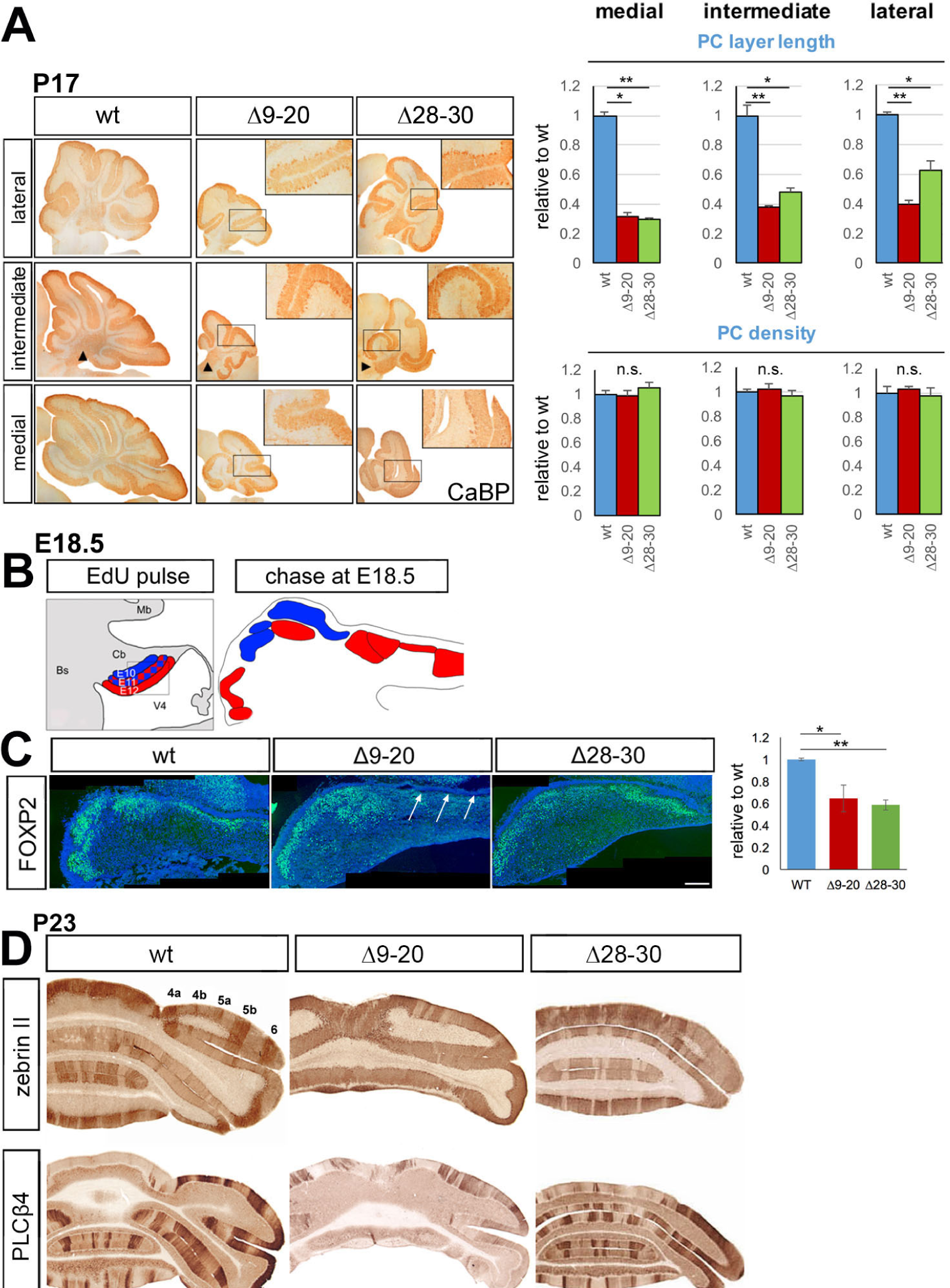
Although PC numbers are reduced in both mutants, the Δ9-20 selectively lacks late born Purkinje cells. **A:** *left,* immunohistochemistry of P17 sagittal sections stained for the PC- specific marker calbindin (CaBP). Cerebellar hypoplasia is evident in both mutants. *Right*, morphometric analysis results point to a severe decrease in PC layer perimeter in both mutants, more pronounced in the cerebellar vermis than the hemispheres; on the contrary, PC numerical density is unchanged. This analysis was conducted using ImageJ (NIH) software and results are plotted as the mean ± SD of biological triplicates. \**p*<0.01; \*\**p*<0.005, Student’s *t-*test. **B:** schematic representation of PC birthdating (EdU pulse at E10, E11 and E12) and PC localization in the cerebellar cortex at E18.5. **C:** FoxP2 immunofluorescence staining of E18.5 frontal sections reveals a severe PC loss in both mutant cerebella, compared to wild type control. Note that only the Δ9-20 displayed a severe reduction of the medial population of PCs corresponding to PCs born at E12.5 (arrows in **B**). The total FoxP2+ PC are counted using the ImageJ (NIH) software, and the result is plotted as the mean ± s.e.m. of biological triplicates (\**p*<0.05; \*\**p*<0.002, Student’s *t*-test). **D**: the results of zebrin II and PLCβ4 immunostaining of P23 frontal cerebellar sections reveal a selective depletion of late-born PCs (PLCβ4+) in Δ9-20 mutants. Conversely, the Δ28-30 mutant cerebellum is smaller but features a balanced representation of early- (zebrin II+) and late-born (PLCβ4+) PCs. Size bar: 200DμM (**C**).

### Allelic *Zfp423* deletions have divergent effects on PC progenitor maintenance and differentiation

To try to clarify the events leading to PC depletion in *Zfp423* mutants, immunofluorescence was used to analyze markers of mitotic (OLIG2+) and postmitotic (CORL2+, LHX5+) PC precursors at E11.5 and E12.5, when PC neurogenesis peaks (Minaki et al., 2008; Seto et al., 2014; Zhao et al., 2007). Our results indicate that at E12.5, the domain containing mitotic (OLIG2+) PC progenitors (**Fig. 4A)**) is significantly reduced in the Δ9-20 cerebellar primordium, while it is expanded in the Δ28-30 primordium. One day earlier, the OLIG2 domain is equally reduced in both mutants (not shown). This is in agreement with the fact that the Δ9-20 mutant selectively lacks late born PCs, while the C-terminal mutant cerebellum, albeit hypoplastic, contains a normal balance of early and late born PCs.

**Figure 4.**
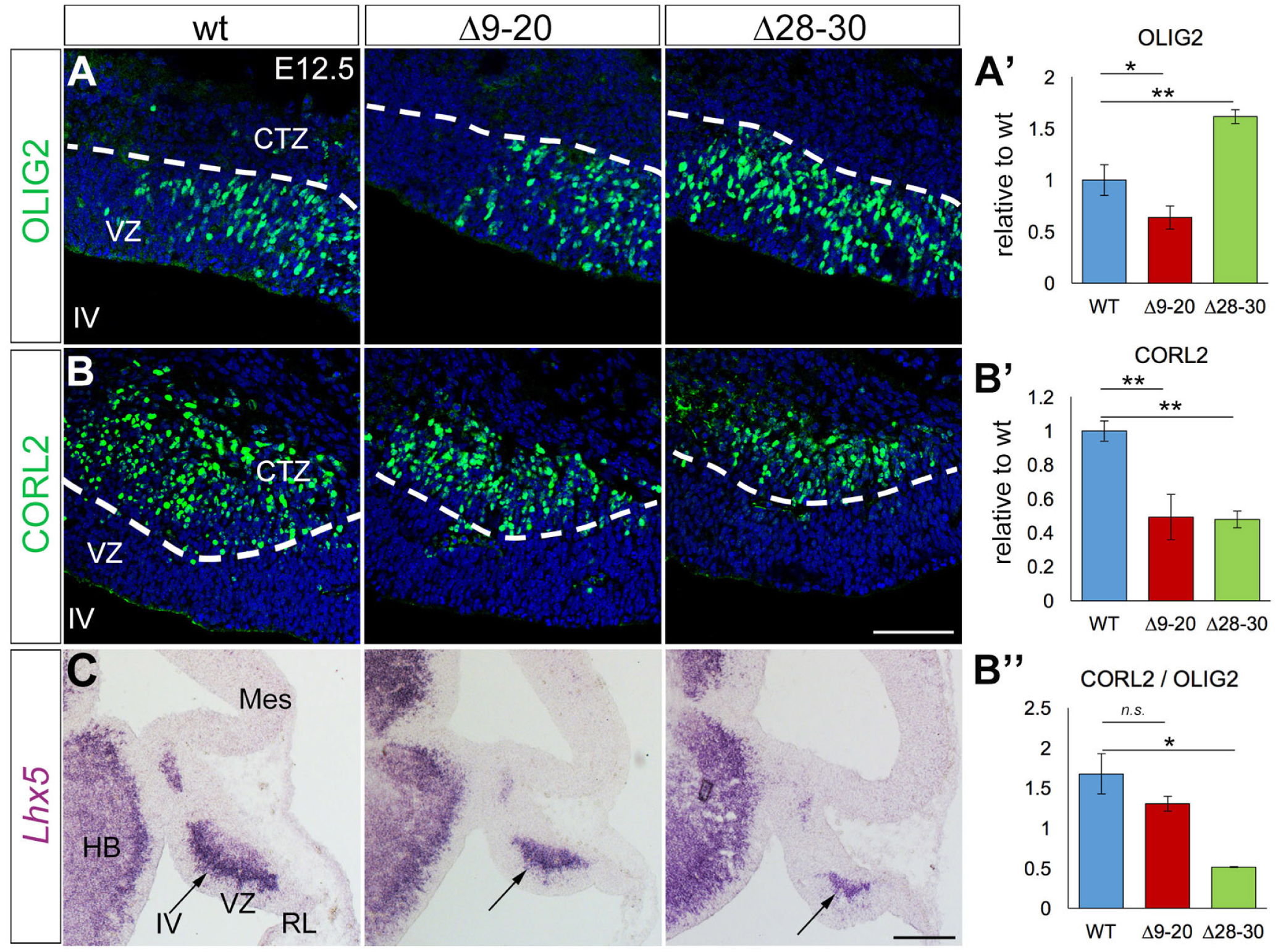
In the cerebellar primordium ZFP423 controls the proliferation and differentiation of PC progenitors. Immunofluorescence staining (**A, B**) and in situ hybridization (**C**) of cerebellar sagittal sections at E12.5 (rostral to the left). **A**: the number of OLIG2+ proliferating PC progenitors is decreased in the Δ9-20 and increased in the Δ28-30, with respect to the wild type control (quantification in **A’**; \**p*<0.05; \*\**p*<0.01, Welch *t*-test). **B**: CORL2+ postmitotic PC precursors are strongly reduced in both mutants (**B**, and quantification in the **B’**; \*\**p*<0.01, Student’s *t*-test), while the CORL2/OLIG2 ratio (**B’’**) is significantly altered only in the Δ28-30 mutants (quantification in the graph; \**p*<0.05, Student’s *t*-test). **C**: *Lhx5*, another marker of early PC differentiation, is downregulated in both mutants, compared with wild type controls. Size bar: 40 μM (**A, B**); 200μM (**C**). All the results are plotted as the mean ± s.e.m. of biological triplicates.

Next, we investigated the role of ZFP423 in the context of PC differentiation. The expression level of CORL2 (**Fig. 4B)**), a PC progenitor differentiation marker, is reduced in the Δ9-20, reflecting the observed decline in the number of OLIG2+ mitotic cells (**Fig.4A**, quantification in **A’**). At the same stage, CORL2 levels (**Fig. 4B)**, quantification in **B’**) are decreased in the Δ28-30 cerebellum, despite the abundance of OLIG2+ cells; this suggests a severe defect/delay in PC progenitor differentiation, such that the CORL2/OLIG2 ratio (**Fig. 4B)’’**) is profoundly decreased in this mutant. The *Lhx5* transcript distribution, analyzed by *in situ hybridization* (**Fig. 4C)**), parallels and supports the described CORL2 results.

As shown, the two allelic mutations have different effects on the ratio of differentiating to proliferating progenitors. Mutations affecting mitotic spindle stability switch the mode of cell division from symmetric/proliferative to asymmetric/neurogenic (Fish et al., 2006), arresting the expansion of the stem cell pool. It is well established that the transition from proliferative to neurogenic cell division of apical radial glial progenitors correlates with a change in the M-phase cleavage plane from vertical to oblique/horizontal, and/or with an unequal inheritance of the apical adherens junction (the “cadherin belt”) in the two daughter cells (Kosodo et al., 2004). Through its dynamic interaction with the centrosomal protein CEP290 (Chaki et al., 2012), itself a Joubert syndrome gene, or via its role as a transcriptional regulator, ZFP423 may affect the stability/orientation of the mitotic spindle.

To evaluate the effect of *Zfp423* mutations on spindle orientation and the type of cell division, we analyzed anaphase figures in the E11.5 cerebellar VZ, in both mutants and controls (representative images in **Fig. 5A))**. Our assessment of symmetric (arrows) vs asymmetric cell divisions (arrowhead) was based on (*i*) expression of the M-phase marker PHH3, (*ii*) the orientation of the plane of cell division with respect to the apical surface, and (*iii*) the equal or unequal inheritance of the adherens junction marker pancadherin (Wilsch-Brauninger et al., 2012). Our results, obtained by analyzing at least three independent embryos per genotype, are shown in **Fig. 5B)**; our data indicate that in the Δ9-20 cerebellar VZ (*n*=60 mitoses tested), the number of asymmetric cleavage planes is significantly increased compared to the wt (*n*=99) (\*\**p*<0.01, Fisher’s exact test), consistent with a shift from symmetric-proliferative to asymmetric-neurogenic cell divisions in this mutant. Conversely, Δ28-30 mitoses (*n*=89) feature a shift towards an excess of symmetric cell cleavages compared to the wt VZ (*p*=0.056, Fisher’s exact test). An comparison of symmetric-to-asymmetric ratios in Δ9-20 vs. Δ28-30 also reveals a strongly significant difference (\*\*\**p*<0.0001, Fisher’s exact test).

**Figure 5.**
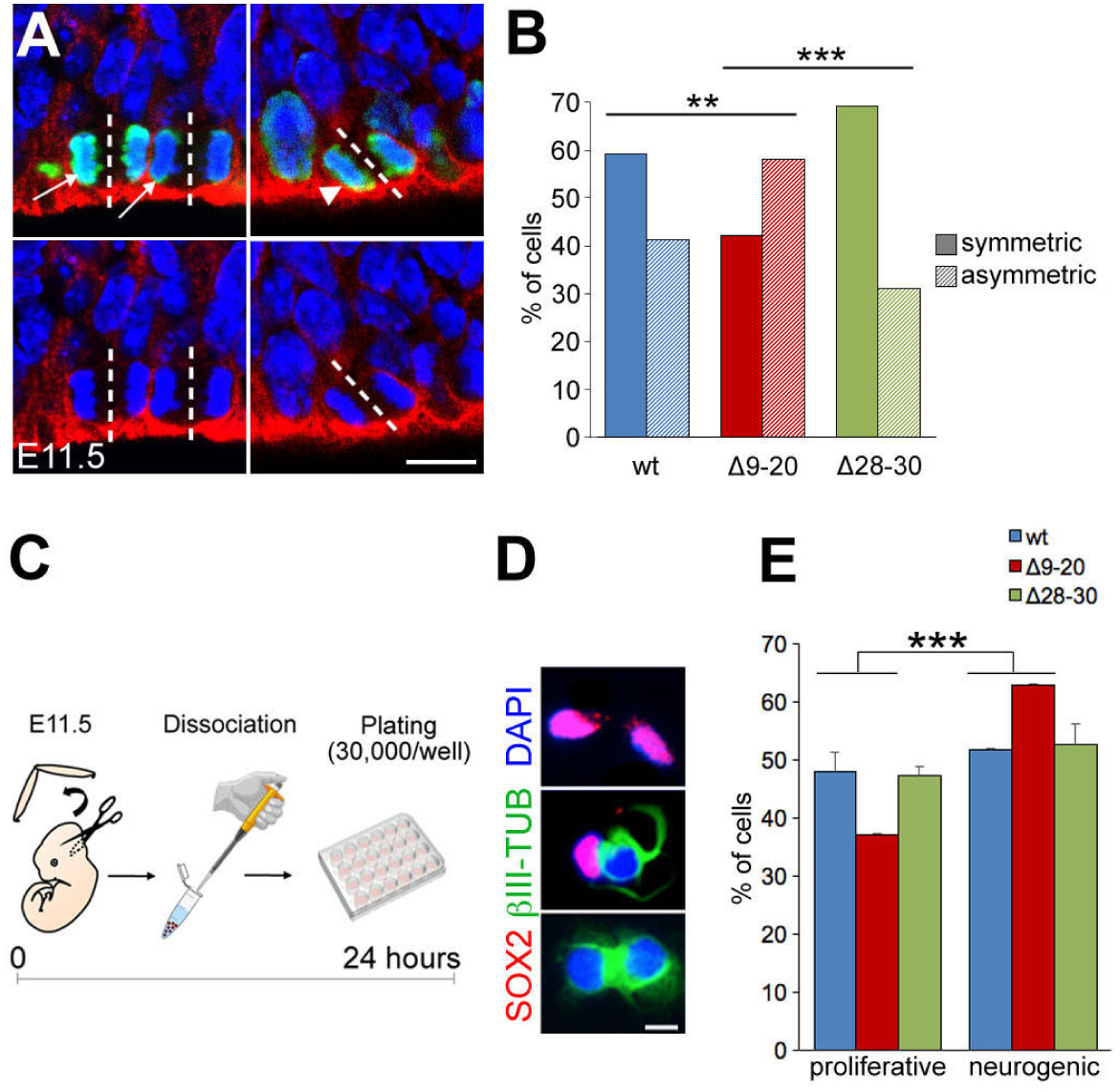
The Δ9-20 deletion causes a premature switch from proliferative to neurogenic progenitor divisions in the cerebellar VZ. **A:** immunofluorescence staining of the E11.5 cerebellar VZ shows the distribution of PHH3, an M-phase marker, and pancadherin (pancdh), decorating the apical adhesion complex. Pancadherin is equally or unequally inherited by the daughter cell during symmetric (proliferative, arrows) or asymmetric (neurogenic, arrowhead) cell divisions, respectively. **B**: quantification of symmetric and asymmetric cell divisions in Δ9-20 and Δ28-30 mutants vs. controls. \*\**p*<0.01; \*\*\*\**p*<0.0001, Fisher’s exact test. **C**: flowchart of the cell pair assay protocol (see Materials and Methods). **D**: sample outcomes of symmetric (PP panel and NN panels, respectively) and asymmetric (PN panel) cell divisions in the cerebellar VZ. Sox2+/Sox2+, proliferative division; βIII-Tub+/βIII-Tub+, self-consuming neurogenic division; Sox2+/βIII-Tub+, asymmentric neurogenic division. **E**: quantification of the percentages of proliferative and neurogenic daughter-cell pairs divisions. The results are plotted as the mean ± SD of biological triplicates \*\*\*\**p*<0.0001, Fisher’s exact test. Size bars: 10μM (**A**); 5μM (**D**).

To determine whether the observed changes in spindle orientation correlate with an overall switch from proliferative to neurogenic cell division, we conducted a clonal cell-pair assay (procedure sketched in **Fig. 5C)**) (Bultje et al., 2009; Li et al., 2003) on primary cultures of cerebellar VZ progenitors. Briefly, E11.5 cerebellar primordia were dissociated and neural progenitor cells (NPCs) plated at clonal dilutions. Cells were fixed 24 hours later and immunostained for SOX2 (undifferentiated progenitors, P) and neural specific β-tubulin III (differentiating neurons, N). Daughter cell pairs were subdivided in two categories: PP, and (NP + NN) (sample mitoses in **Fig. 5D)**). Our results (**Fig. 5E)**) indicate that Δ9-20 NPCs (*n*=852) display a cumulative increase in the ratio of neurogenic (NP+NN) to proliferative (PP) divisions, compared to wt and Δ28-30 (*n*=1339 and 1021, respectively) (\*\*\**p*<0.0001, Fisher’s exact test). Thus, both *in vivo* and *in vitro* data point to a role for the domain spanning ZFs 9-20 in maintaining a pool of undifferentiated progenitors in the cerebellar VZ.

### The DNA damage response is impaired in *Zfp423-*mutant cerebellar VZ progenitors

In an attempt to explain the overall decrease in PC numbers observed in both mutants, we analyzed cell cycle progression of VZ progenitors at E12.5 in three independent embryos per genotype. In particular, we examined the transition between S and M phase, which encompasses the G_2_-M cell cycle checkpoint. To this end, pregnant dams were given a single EdU pulse 30 min prior to sacrifice. Embryonic day 12.5 cerebellar sections were stained for EdU and the M-phase marker PHH3. Our results (**Fig. 6A)**) indicate that although the number of S phase progenitors is not diminished in our mutants, both exhibit a significant increase (about 40%) in the ratio of S- to M-phase progenitors, suggesting that cell cycle progression is arrested, at least in some cells, during this transition (\**p*<0.01 and \*\**p*<0.001, respectively; Fisher’s exact test). When we pulsed pregnant dams 3 hours before sacrifice, in the mutant progeny, all M phase cells (PHH3+) had incorporated EdU, but their fraction was again sharply reduced (not shown). Considering that at E12.5 (G_2_+M) duration equals 2 hours, this suggests that in *Zfp423* mutants some cells progress through the S-M phase at a normal pace, while other cells are stalled. To further confirm this observation, we stained sections for PHH3 and looked at the ratio of dotted PHH3+ (G_2_) to solid PHH3+ (M phase) signal. Again, our results clearly indicate that both the Δ9-20 and Δ28-30 cerebellar primordia feature an increased ratio of G_2_ to M progenitors (\**p*<0.01 and \*\**p<*0.001, respectively; Fisher’s exact test) (**Fig. 6B)**).

**Figure 6.**
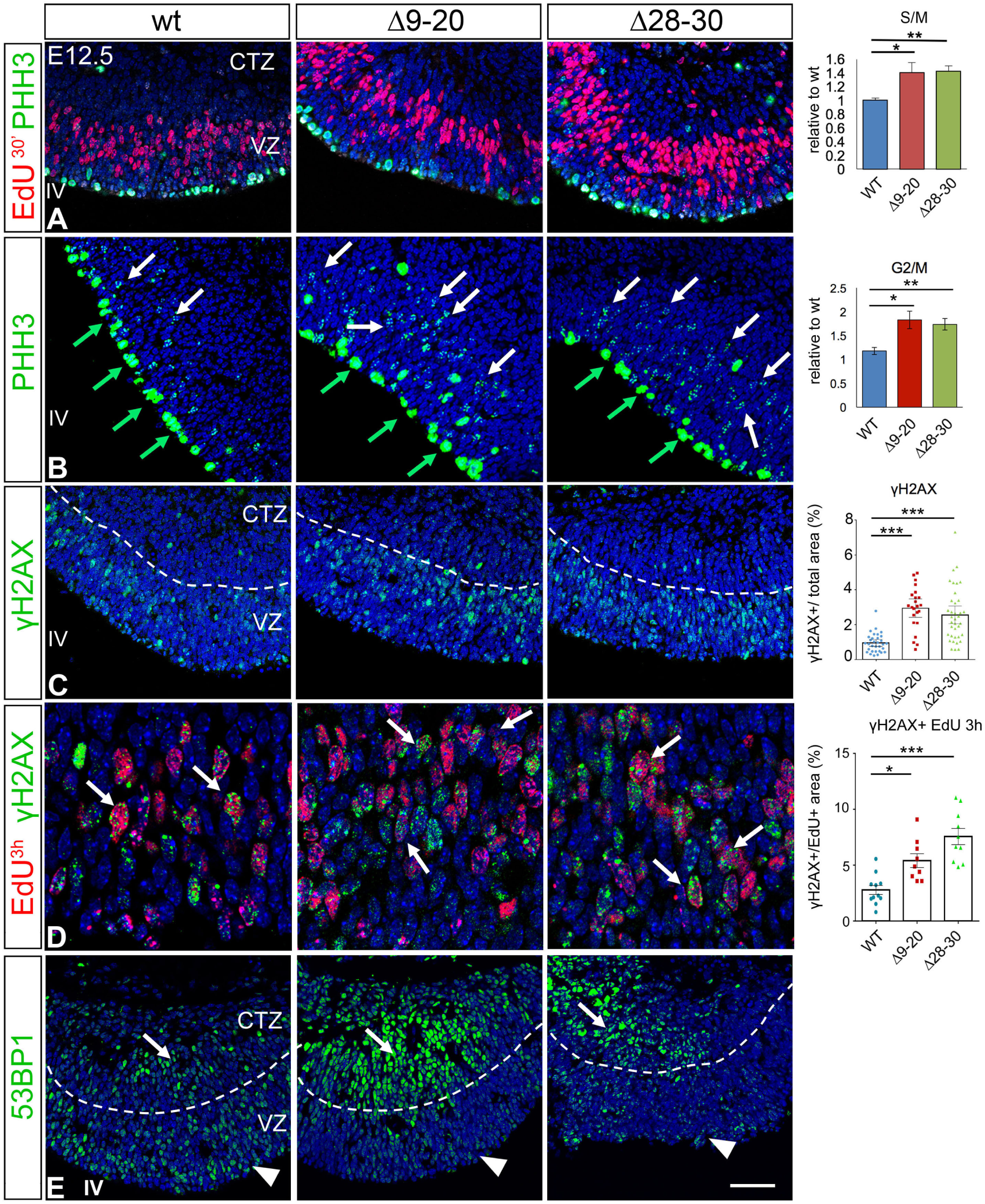
Stall of cell cycle progression and increase in DNA damage signaling in the VZ of ZFP423 mutants. **A-E:** Sagittal sections of the cerebellar primordium at E12.5 stained for markers of specific cell cycle stages and DNA damage signaling (rostral to the left). **A:** PHH3 and EdU (single 30’ pulse) label M- and S-phase progenitors, respectively. The graph on the right shows that the ratio between S and M phases is increased in the mutants (\**p*<0.01 and \*\**p*<0.001, Fisher’s exact test). **B:** dotted and full PHH3 label G2 and M phase progenitors, respectively. The graph on the right shows that the ratio between G2 and M phases is increased in the mutants (\**p*<0.01 and \*\**p*<0.001, Fisher’s exact test). **C:** the density of γH2AX staining, a marker of double strand breaks, is significantly increased in the cerebellar VZ of both mutants. ( \*\*\**p*<0.0001, one-way ANOVA, Kruskal-Wallis test, Dunn’s multiple comparison test); **D:** colocalization of γH2AX with EdU (red), shown by arrows, after a 3hr single pulse is strongly upregulated in both mutants. (\**p*<0.02; \*\*\**p*<0.0001, one-way ANOVA, Kruskal-Wallis test, Dunn’s multiple comparison test). **E**, in both mutants, 53BP1 expression (green) is significantly upregulated in the cortical transitory zone (arrows) and downregulated in the vz (arrowheads). Size bars: **A-C**,**E**: 40 μM; **D**: 15 μM. All the results are plotted as the mean ± s.e.m of biological triplicates. IV: 4th ventricle; VZ: ventricular zone; CTZ: cortical transitory zone.

ZNF423 has been linked to the DNA damage response (Chaki et al., 2012). DNA damage repair is tightly connected to the regulation of cell cycle progression (reviewed in Branzei and Foiani, 2008). Single-strand gaps or nicks and double strand breaks (DSBs) often occur during S phase and trigger repair by nonhomologous end joining (NHEJ) and/or homologous recombination (HR) in mitotic cells. In normal neural tube development, the rapid pace of cell division set by neural progenitors likely exposes them to a high degree of endogenous DNA damage (stalled replication forks leading to single and double strand breaks), requiring repair prior to cell division (reviewed in McKinnon, 2013). The finding of mutant progenitors stalled during their G_2_-M phase transition prompted us to ask whether DNA damage signalling is increased in vivo in *Zfp423* mutants. To test this hypothesis, we immunostained E12.5 cerebellar sections for the DNA damage marker phosphorylated histone 2A family member X (γH2AX) (Celeste et al., 2002). Our results (**Fig. 6C)**) point to a sharp increase in DNA damage signaling in the cerebellar VZ of both mutants (\*\*\**p*<0.0001, Kruskal-Wallis test). By immunofluorescence, we assessed the percentage of colocalization between EdU^3h^+ cells (S-M) and γH2AX, and observed a significant increase in colocalization in both mutants (\**p*<0.02; \*\*\**p*<0.0001, Kruskal-Wallis test). Finally (**Fig. 6E)**), we observed a qualitative and sharp increase in the expression of 53BP1, a protein involved in NHEJ in response to DSBs (reviewed in Panier and Boulton, 2014), in postmitotic precursors (arrows in **6E**), indicating that mitotic progenitors carry over DNA damage upon cell cycle exit, likely due to a reduced efficiency of the DDR in the cerebellar VZ, and resort to NHEJ for DNA repair. Interestingly, the expression of 53BP1 in the VZ is downregulated in *Zfp423* mutants, particularly in the Δ9-20, compared to wt (arrowheads in **6E**), possibly suggesting a decreased recruitment of 53BP1 on DNA damage foci in dividing cells.

Taken together, these results strongly suggest that in both mutants the frequency of double-strand breaks is significantly increased throughout the cell cycle, likely due to inefficient repair, causing an equally sharp increase in residual DNA damage in differentiating neurons.

## DISCUSSION

The key role played by *Zfp423 / ZNF423* in cerebellar morphogenesis has been addressed through the analysis of mouse models and human patients alike. Studies conducted by others on mouse lines carrying engineered or spontaneous null mutations of *Zfp423* have elucidated the importance of this gene in cerebellar organogenesis (Alcaraz et al., 2006; Cheng et al., 2007; Warming et al., 2006). Moreover, through a combined homozygosity mapping and exome analysis approach, mutations of *ZNF423*, the human ortholog of murine *Zfp423*, have been identified in patients featuring Joubert syndrome and nephronophthisis, classifying *ZNF423* as a human ciliopathy gene (Chaki et al., 2012). All JS proteins identified prior to this discovery are components of the primary cilium or basal body. However, the evidence supporting a centrosomal localization of ZFP423 is still inconclusive. In our hands, ZFP423 features a clear-cut nuclear localization, in agreement with its established role as a TF (Hata et al., 2000; Ku et al., 2006; Ku et al., 2003; Masserdotti et al., 2010; Tsai and Reed, 1997; Tsai and Reed, 1998).

The role of the cilium in GC development is well established (Chizhikov et al., 2007) and relates to this organelle’s function in the context of SHH signaling and other regulatory pathways. Other authors have convincingly demonstrated that ZFP423 regulates the proliferative response of GC progenitors to SHH (Hong and Hamilton, 2016). However, our work also implicates ZFP423 in the control of PC development, a process that is independent of SHH signaling and the role, if any, of the primary cilium in this process has not been addressed to date. Incidentally, the overall number of GCs in the mouse cerebellum is determined by the number of immature PCs releasing SHH between the late fetal period and postnatal day 15, concatenating the two processes (Dahmane and Ruiz-i-Altaba, 1999; Wallace, 1999; Wechsler-Reya and Scott, 1999). In humans, JS can be clearly diagnosed as a cerebellar malformation in utero by magnetic resonance imaging (Saleem et al., 2011u887), while the bulk of GC clonal expansion occurs after birth in our species (Kiessling et al., 2014), as in mice. JS research has dealt more with GCs than with PCs in both human patients and murine models. In an attempt to fill this gap we have focused on early stages of PC development.

### The Δ9-20 mutation affects the maintenance of mitotic PC progenitors

The two mouse lines analyzed in this work carry distinct, functionally characterized ZFP423 protein domains. The two mutations, while both giving rise to stable nuclear proteins, do not produce any detectable gain of function: in fact, heterozygotes are morphologically and functionally normal, while homozygotes have cerebellar hypoplasia and are severely ataxic (not shown). Our findings uncover some specific roles in PC development for each of the deleted ZFP423 protein domains. The deletion of ZF’s 9-20 causes a precocious depletion of late-born PCs (PLCβ4+), whereas early-born PCs (zebrinII+) are unaffected. As a result, the alternating pattern of zebrinII+ and - stripes is lost in the Δ9-20 cerebellum, whereas it is maintained in the Δ28-30 cerebellum, despite its small size. Accordingly, at E11.5, the Δ9-20 cerebellar VZ displays changes in M-phase progenitor spindle orientation and an unequal inheritance of the cadherin+ adherens junction (Paridaen et al., 2013). Notably, at this early stage, the wt VZ features a predominance of proliferative divisions. Likewise, in the cell pair assay (Bultje et al., 2009), this change translates into an increase in the ratio of neurogenic to proliferative cell divisions in Δ9-20 NEPs but not in Δ28-30 NEPs. Incidentally, because ZFP423 signal peaks in M-phase VZ progenitors and appears to colocalize with nuclear membrane components (lamin B) and to segregate from compacted chromatin, the possibility of a transcription-independent role of this protein in mitotic spindle stabilization cannot be excluded, and should be addressed experimentally.

### Zinc fingers 28-30 of ZFP423 regulate PC progenitor differentiation

In contrast with the observations made in the Δ9-20 primordium, the allelic mutant Δ28-30 shows no decline of OLIG2+ progenitors, in keeping with the fact that late-born PCs are present in this strain, and yet features a severe defect/delay in their differentiation into CORL2+ and *Lhx5+* postmitotic precursors. The C-terminal domain of ZFP423 mediates the interaction of this protein with EBF transcription factors (Hata et al., 2000; Masserdotti et al., 2010; Tsai and Reed, 1997; Tsai and Reed, 1998), which play a key role in coupling cell cycle exit with the onset of neuronal migration and differentiation in neural tube development (Garcia-Dominguez et al., 2003). Two EBF transcription factors, EBF2 and EBF3 (Malgaretti et al., 1997; Wang et al., 1997), have been implicated in mouse and human cerebellar development, respectively (Chao et al., 2017; Chung et al., 2008; Croci et al., 2006; Harms et al., 2017; Sleven et al., 2017). EBF2, in particular, affects various aspects of PC development, namely the migration and survival of a late-born PC subset. In the Δ28-30, PC progenitor differentiation may be delayed or impaired due to the lack of ZFP423-EBF interaction. However, the Δ9-20 deletion does not affect the interaction of ZFP423 with EBF TF’s, perhaps sequestering ZFP423 into an inactive heterodimer and shifting the balance from progenitor maintenance to cell cycle exit.

Although ZFP423 is expressed at robust levels in VZ progenitors, suggesting a cell-autonomous role of this protein, it is also found in rhombic lip and roof plate derivatives. While there is no known effect of RL progenitors on PC development prior to PC migration, extracellular factors secreted by the roof plate and choroid plexus may possibly contribute to the alterations observed in our mutants. Conditional inactivation of *Zfp423* in different lineages should provide answers to these open questions.

### Both ZFP423 deletions cause cell cycle progression arrest in a subset of G2 cells

In this work we studied cell cycle progression in our mutant lines by analyzing the expression of global and cell-type specific markers. DNA replication was assessed by a single 30’ EdU pulse at E12.5. At this stage, the ratio of EdU incorporation to DAPI staining was unchanged (not shown), in keeping with previous reports (Alcaraz et al., 2006). However, the S/M and G2/M ratios were significantly increased in both mutants, suggesting that some cells are delayed in their progression from S to M phase. Given the role of ZFP423 in the DDR (Chaki et al., 2012), we speculated that ZFP423 mutation might cause a subset of progenitors to stall during the same interval. In particular, the delay in G_2_/M progression strongly suggests that ZFP423, which is expressed at very low levels in G_1_ progenitors (Suppl. Fig. 1A), is involved in permitting transition through the G2/M cell cycle checkpoint. This evidence prompted an in vivo analysis of DNA damage in PC progenitors.

### Both ZFP423 deletions cause an increase in DNA damage signaling in the cerebellar primordium

While each deletion affects separate aspects of PC progenitor development, both mutants feature a massive increase of DNA damage signaling in the cerebellar VZ. The 9-20 domain of ZFP423 overlaps almost entirely with the region of the protein known to interact with the DNA ds-damage sensor PARP1 (poly-ADP ribosyl polymerase 1) (Ku et al., 2003). PARP1 decreases the affinity of the Ku70-Ku80 heterodimer for DS breaks, shifting the balance of DNA repair from NHEJ to HR (Hochegger et al., 2006). In the context of the DDR, HR occurs, by definition, after DNA replication, to recruit a homologous chromatid (usually a sister chromatid) as a template for DNA repair. In order for HR-mediated repair to occur, cell cycle progression must be arrested so as to delay chromatin compaction, which would make the search for homology difficult. Our evidence suggests that DNA damage signaling is significantly increased in S-M progenitors. Conceivably, in the Δ9-20 cerebellum, which loses the PARP1 interaction domain, a defective recruitment of PARP1 could prevent the assembly of a functional DNA repair complex acting between S and M, presumably by HR. While ZFs 9-20 contain the PARP1 interaction domain and are immediately adjacent to the CEP290 interaction site (Chaki et al., 2012), no clues are available to date as regards the function of ZF’s 28-30 in the context of DNA repair. We hypothesize that the integrity of the ZFP423 C-terminus is necessary to assemble a functional DNA repair complex with as yet unidentified co-factors. Possibly because of the diminished efficiency of HR in mitotic cells, both mutants sharply upregulate the DNA end resection inhibitor 53BP1 (Escribano-Diaz et al., 2013) in postmitotic cerebellar precursors, which are substantially dependent on NHEJ for DNA repair.

### Ciliopathy genes, the DNA damage response and Joubert syndrome

The finding of a highly significant increase of endogenous DNA damage in cycling VZ progenitors further supports the emerging notion that several ciliopathy genes play important extra-ciliary roles in the nucleus, participating in the DDR. This has been shown elegantly in renal ciliopathies: both *CEP290* and *NEK8* mutations (Choi et al., 2013; Slaats et al., 2015) lead to an accumulation of DNA damage due to disturbed replication forks. Furthermore, increased DNA damage signaling has been scored in *CEP164*-, *ZNF423*-, and *SDCCAG8*-associated nephronophthisis (Airik et al., 2014; Chaki et al., 2012). This study raises the possibility that cerebellar hypoplasia in JS, and in other cerebellar malformations, may stem at least partially from defective DNA repair. While in this study we have focused on PC development, an increased DNA damage may in fact contribute to the altered response to SHH observed by others in fast-dividing granule cell precursors of the EGL.

In conclusion, the mounting evidence of defective repair of endogenous DNA damage in fast-cycling cerebellar VZ progenitors furthers our understanding of Joubert syndrome, and may have far reaching implications in our overall approach to this elusive disorder.

## ACKNOWLEDGEMENTS

Image analysis was carried out in ALEMBIC, an advanced microscopy laboratory established by the San Raffaele Scientific Institute and Vita-Salute San Raffaele University. We thank F. Bani and G. Bergamini for their contribution to this project. This work was mostly funded by the Italian Telethon Foundation through grant GGP13146 to GGC; a partial contribution was provided by Ministero della Salute through grant “Ricerca Finalizzata 2011” PE-2011-02347716 to OC. RH was supported by an award from the Canadian Institutes of Health Research. SW and LT were supported by the NIH Intramural Research Program, Center for Cancer Research, National Cancer Institute.

## SUPPLEMENTAL FIGURE LEGENDS

**Supplemental figure 1.**
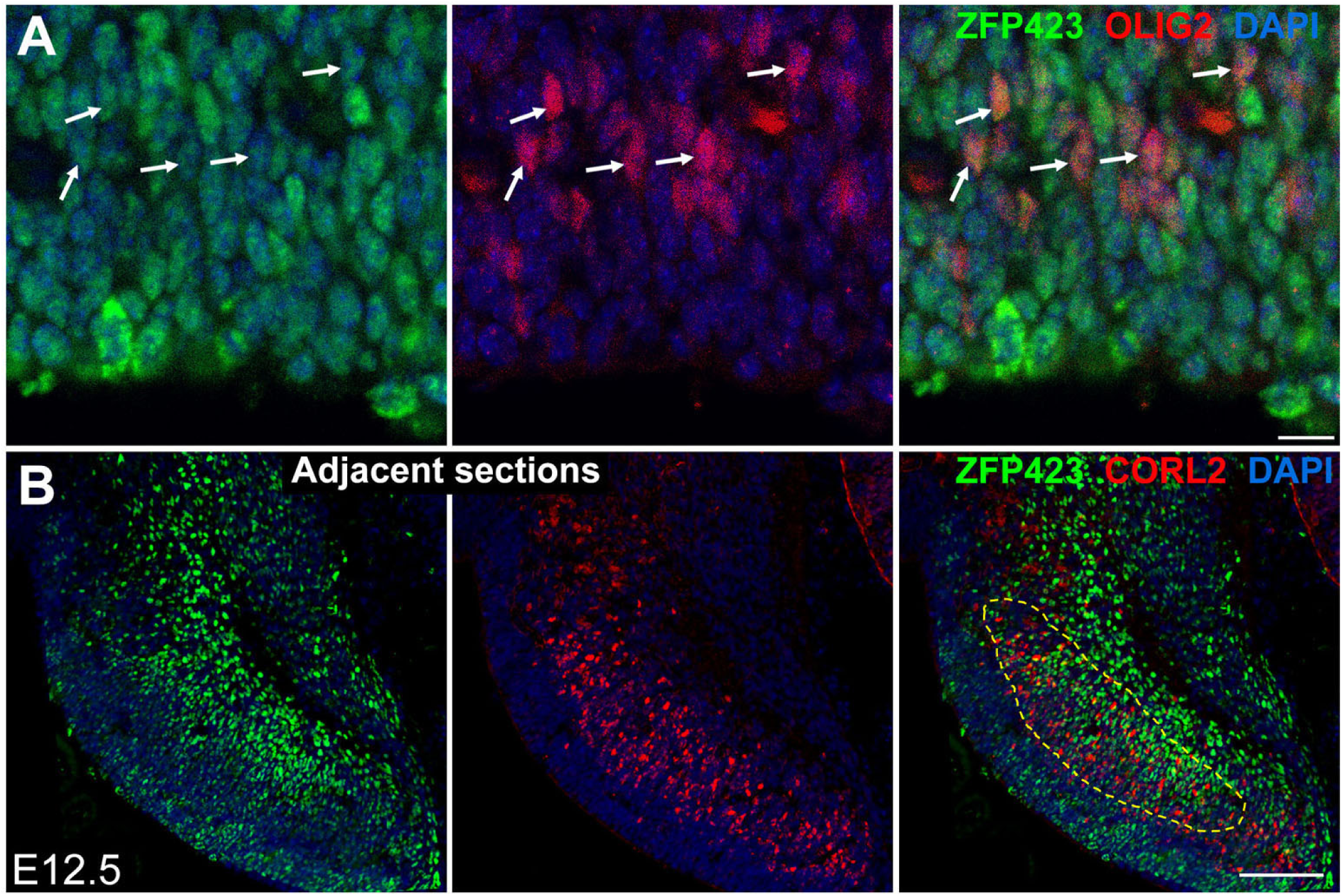
Immunofluorescence staining of cerebellar sagittal sections at E12.5. ZFP423 is expressed in some OLIG2+ cells (**A**, arrows) and overlaps partially with CORL2+ domain (**B**). Size bars: **A**, 20μM; **B**, 100μM.

**Supplemental figure 2.**
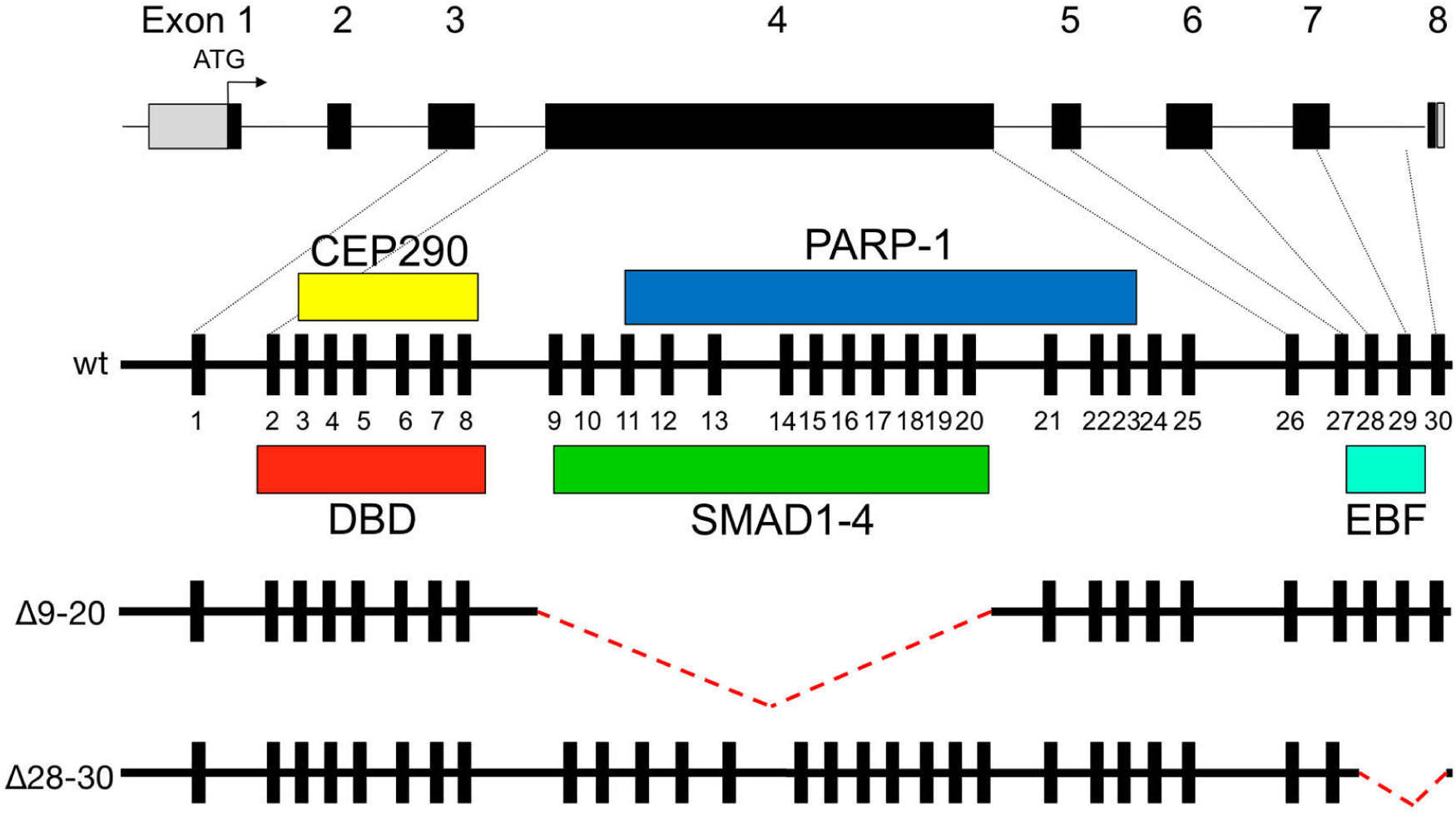
ZFP423 / ZNF423 has been characterized as a nuclear scaffold protein mediating multiple protein-protein interactions. Briefly: CEP290 is a centrosomal protein implicated in centrosome and cilium development; PARP1 is a chromatin-modifying enzyme that triggers ADP-ribosylation of other nuclear proteins and is involved in DNA damage repair; SMAD proteins are nuclear transducers of BMP signaling; EBFs are transcription factors implicated in neuronal differentiation, migration and survival. See text for references.

